# Spatial sorting and accelerated larval development in a range expanding fiddler crab

**DOI:** 10.64898/2025.12.16.694704

**Authors:** Jordanna Barley, Eric Sanford, Brian S Cheng

## Abstract

Although climate change has driven the widespread occurrence of species’ range shifts, the ecological and evolutionary processes that generate these patterns remain poorly understood. Here, we investigate spatial sorting and countergradient selection as evolutionary mechanisms contributing to range expansion. We focus on the rapid range expansion of a fiddler crab (*Minuca pugnax*) in the Gulf of Maine, an ocean warming hotspot. We use common garden experiments to quantify change in pelagic larval duration (PLD), a key determinant of dispersal ability, across edge and interior core populations, and between timepoints separated by 19 years. Consistent with spatial sorting, larvae from poleward, edge populations had shorter PLDs relative to equatorward, core populations, which is more dispersive in this system because it is characterized by equatorward flows. Compared to 19 years prior, PLDs were significantly shorter, consistent with genetic change over time. Analysis of sea surface temperatures reveal that the larval development window in the edge populations has doubled over the past twenty years, suggesting the combined influence of ocean warming, trait differentiation over time, and spatial sorting in promoting range expansion. While spatial sorting is known to aid in terrestrial range expansions, our results are the first to suggest that this process might be important in the coastal oceans.

## Introduction

Anthropogenic climate change is causing a global reorganization of biodiversity, including shifts in species’ ranges that can manifest as local extirpation and range contraction at lower latitudes and expansion towards the poles (Parmesan 2006; Chen et al. 2011; Pecl et al. 2017). The ubiquity of range shifts across taxa and ecosystems has been regarded as a universal response to climate change (Walther et al. 2002; Parmesan and Yohe 2003; Daufresne et al. 2009; Lenoir and Svenning 2015; Rubenstein et al. 2023). Range shifts may be driven by multiple factors, including direct dispersal by adults, overwintering survival, and transport and recruitment of dispersive young (Pinsky et al. 2020; Rubenstein et al. 2023). However, there remains considerable variability in the speed and direction of range shifts, with some species exhibiting altered distributions that are seemingly inconsistent with the predicted effects of climate change (e.g., equatorward or towards shallower waters; Poloczanska et al. 2013; Fuchs et al. 2020). Thus, the mechanisms underlying range shifts are not well understood and exhibit considerable complexity.

Although many processes can influence range shifts, dispersal capacity is central to the colonization of new habitats. Traits that promote dispersal often accumulate at a species’ range edge due to spatial sorting, an evolutionary process that operates outside of natural selection (Shine et al. 2011; Miller et al. 2020). Spatial sorting occurs because high dispersing individuals are more likely to accumulate at a species’ range edge simply due to their ability to disperse farther. Individuals at the range edge may also undergo assortative mating, producing offspring that also have high dispersal capacity (i.e., the ‘Olympic village effect’; Phillips et al. 2010). For example, dispersive plant traits such as smaller seeds and higher germination rates are found in expanding range edge populations (Cwynar and MacDonald 1987). In bush crickets, longer-winged morphs are more likely to be found at the range edge because they can fly farther and disperse more widely (Simmons and Thomas 2004). Over time, as a species continues to expand poleward, range-edge populations are incorporated into the range core and should be less influenced by spatial sorting. Therefore, the average dispersal capacity in transient range-edge populations may decline as a range expansion proceeds (Simmons and Thomas 2004). Although traits that increase an individual’s dispersal potential can be correlated with traits that increase an individual’s fitness (Burton et al. 2010; Perkins et al. 2013; Peischl and Gilbert 2020), spatial sorting does not inherently increase survival and reproduction and can result in decreased fitness at the range edge (Shine et al. 2011; Shaw et al. 2023). Despite growing evidence for spatial sorting in terrestrial ecosystems, the role of this evolutionary process in contributing to range expansion is not well resolved, particularly in marine taxa that experience greater climate velocities and appear to be shifting up to 3-11x faster than in terrestrial systems (Parmesan and Yohe 2003; Loarie et al. 2009; Sorte et al. 2010; Chen et al. 2011; Poloczanska et al. 2013).

In marine systems, many invertebrates and fish disperse primarily or exclusively as small pelagic larvae (Thorson 1950). The time spent in this life stage is termed the pelagic larval duration (PLD), which can have a large influence on a species’ dispersal capacity, geographic range, and capacity to range shift (Siegel et al. 2003). Because temperature primarily accelerates metabolic processes, PLD shortens as temperature increases, leading to the hypothesis that climate warming will decrease dispersal potential and population connectivity (Houde 1989; O’Connor et al. 2007; Munday et al. 2009), potentially reducing species’ capacities for range expansion.

Range shifts may also be limited by physical processes. In many coastal ecosystems, ocean currents are dominated by equatorward flows (e.g., California, Western Gulf of Maine, Mid-Atlantic Bight, Benguela, and Humboldt current systems), yet poleward range shifts of species with planktonic larvae are still common within these environments (Audet et al. 2003; Zacherl et al. 2003; Epifanio 2013; Johnson 2014, 2015). For example, the 2014-2016 marine heatwaves in the California Current Large Marine Ecosystem drove the range expansion of 37 marine taxa, 21 of which were benthic animals with planktonic larval dispersal (Sanford et al. 2019). In this context, ocean warming can promote poleward range expansion via at least two mechanisms. First, ocean warming could increase environmental temperatures past a thermal threshold for development, which allows for the completion of the larval stages in habitat beyond a species’ historic range (Sanford et al. 2006; Ling et al. 2008). Second, warming increases developmental rates and shortens organismal PLD. Upon first inspection, such a phenomenon might be expected to decrease dispersal potential and the ability for species to colonize new habitats. However, a shorter larval duration could act in concert with episodic poleward current reversals to promote the poleward dispersal of propagules (Zacherl et al. 2003; Sanford et al. 2019). Indeed, this is consistent with theoretical simulations in advective environments (Byers and Pringle 2006) and is analogous to the “drift paradox” in freshwater stream systems, which seeks to resolve how aquatic taxa can maintain their spatial distribution when living within environments dominated by directional transport (Hershey et al. 1993; Williams and Williams 1993). Thus, in the many marine ecosystems characterized by equatorward flows, short PLDs that are typically thought of as less dispersive might be more dispersive (Sanford et al. 2006). If true, evolutionary theory suggests that this trait may be spatially sorted at the range expansion edge.

Rapid development in poleward marine populations may also be indicative of countergradient variation (CnGV), a pattern arising from local adaptation whereby the genotypic effects on a trait oppose environmental effects (Conover and Present 1990). For example, colder temperatures associated with higher latitudes are expected to decrease growth rates yet it is commonly observed that populations from higher latitudes have greater developmental rates as compared to their low latitude counterparts when reared at the same temperature (reviewed in Conover et al. 2009). This may arise because shortened growing seasons at high latitudes may select for developmental rates that promote overwintering survival and greater lifetime reproductive success in their home environment (Conover and Present 1990; Laugen et al. 2003). Numerous empirical examples exist across ecosystems and taxa showing that countergradient variation arises in range-expanding species (Laugen et al. 2003; Sanford et al. 2006; Carbonell and Stoks 2020; Sefbom et al. 2022). However, countergradient variation is typically evaluated across space (i.e., synchronic comparisons; Kawecki and Ebert 2004), but it is a process that can also operate across time (Garant et al. 2004; Conover et al. 2009). To our knowledge, only one study has evaluated the influence of temporal warming on CnGV, which resulted in a finding opposed to theory; developmental rates in tadpoles increased instead of decreased over a period of warming (Arietta and Skelly 2021). Their approach was to repeat common garden experiments across time, comparing development in the same populations that have undergone environmental change (i.e., allochronic comparisons; Nevo et al. 2012). Importantly, spatial sorting and countergradient variation are not mutually exclusive and may operate in tandem to produce highly dispersive and locally adapted populations at the edge of a range (Van Petegem et al. 2016; Miller et al. 2020).

Here, we evaluate the evolutionary processes that can shape range expansion by examining the mud fiddler crab (*Minuca pugnax,* formerly *Uca pugnax*), a species that is undergoing rapid range expansion in the Gulf of Maine. Our objectives were to 1) investigate the potential for spatial sorting or countergradient variation by contrasting PLD from two edge populations and two range-core populations, 2) quantify metabolic rate of larvae via respirometry, to evaluate a possible mechanism underlying patterns of differing development time, and 3) test whether PLDs of larvae from two edge populations have changed in association with warming ocean temperatures between 2003-04 and 2022. In addition, 4) we used remote-sensed sea surface temperature (SST) data to explore the relationship between environmental data and PLD over this 19-year period. We hypothesized that a clear signal of spatial sorting (i.e., shorter PLD in edge populations), would manifest in comparisons of edge vs. core populations that are subject to large geographical separation and reduced population connectivity. We also hypothesized that larvae with faster larval development would be characterized by greater metabolic rates, consistent with the metabolic compensation hypothesis (Conover et al. 2009). If there was a pattern of countergradient variation, we hypothesized that rapid warming within this system would result in relaxed selection for faster development in larvae, thereby producing greater PLDs in the edge populations over time (i.e., decreased developmental rates).

## Methods

### Study System

The Gulf of Maine (GOM) is one of the fastest warming coastal areas in the world (Pershing et al. 2015; Townsend et al. 2023) and is characterized by an equatorward coastal current at the nearshore where many larval propagules are dispersed (Pettigrew et al. 2005). This coastal current continues south of Cape Cod and into the Mid-Atlantic towards the interior of the mud fiddler crab distribution (Roarty et al. 2020). There are multiple documented poleward range expansions in the GOM (Johnson 2014, 2015; Bell et al. 2015) including the mud fiddler crab. This species disperses using pelagic larvae (Sanford et al. 2006; Johnson 2014), and has a historic distribution from Florida to Cape Cod, MA, and is now expanding into the Gulf of Maine. Mothers hold developing embryos for an incubation period of approximately 2-3 weeks and broadcast larvae into the water column (Brown and Loveland 1985). Sanford et al. (2006) found northern populations of *M. pugnax* had shorter PLDs than those on the south side of Cape Cod, but the pattern varied between consecutive years. In addition, they identified 18°C as the lower thermal threshold that *M. pugnax* requires to complete larval development and metamorphose. Since 2000, there has been a striking increase in the number of days above 18°C in the GOM from an average of less than 40 to more than 80 days (Figure 1), likely promoting the range expansion of *M. pugnax*.

**Figure 1.**
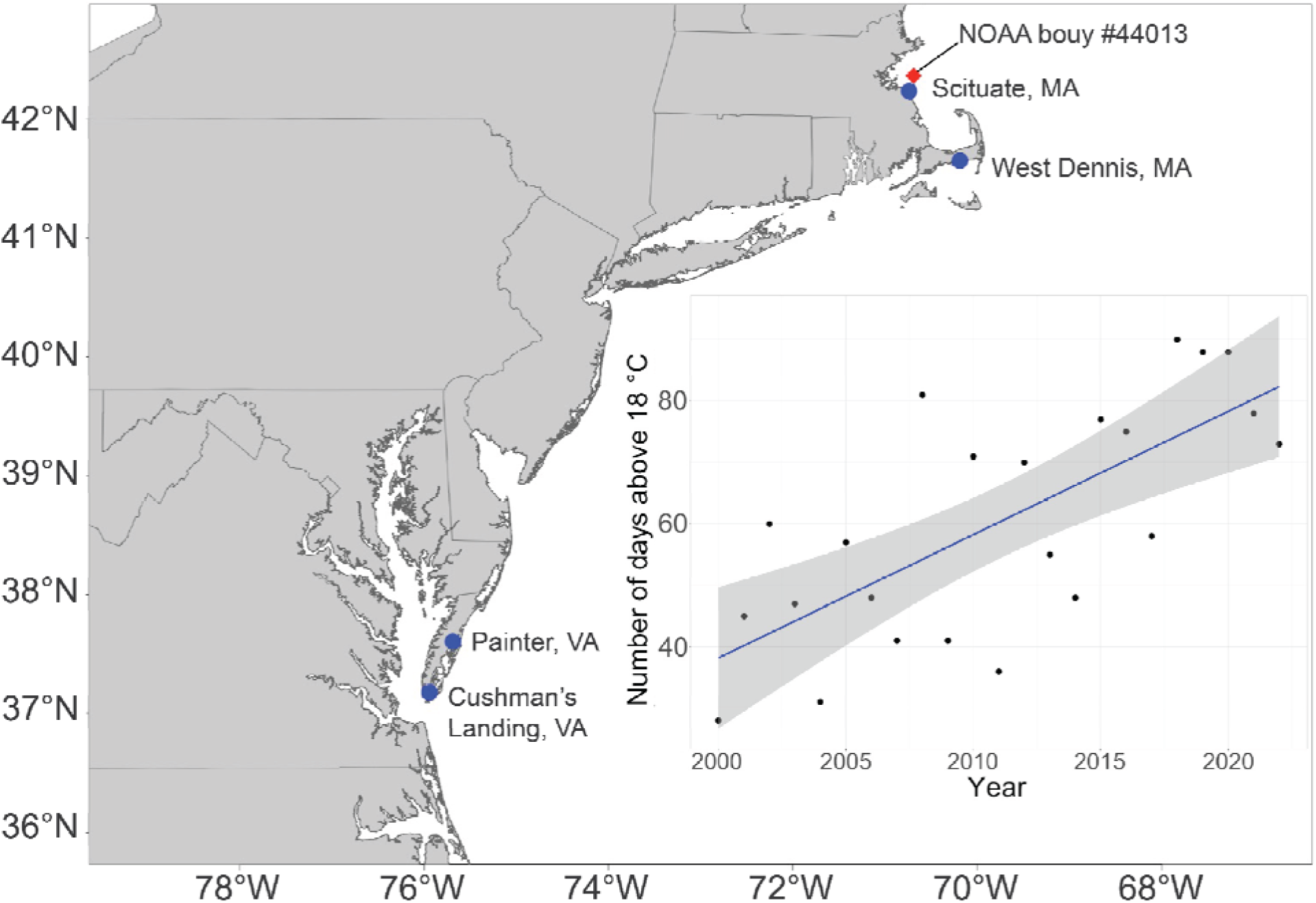
Map of the east coast of the US with blue dots denoting collection sites for *Minuca pugnax* broodstock in the 2022 experiment. The two northern populations were also sampled in the 2003 and 2004 experiments (Sanford et al. 2006) as well as in the 2019 experiments. Inset figure shows the number of days per year with daily average temperature above 18□in the Gulf of Maine (NOAA buoy #44013, location depicted by red diamond). Model fit line is the result of linear regression with 95% confidence interval bands (F_1,22_ = 27.71, p<0.0001).

### Field collection of ovigerous females

To obtain estimates of PLD as a dispersive trait across space and time, we collected ovigerous female *M. pugnax* within a week of the first spring tide in June during 2019 and 2022. Collecting females ahead of the first spring tide of the spawning season maximized the chances of synchronized spawning and is consistent with past experiments (Wheeler 1978; Christy and Stancyk 1982; Sanford et al. 2006). We collected females from a total of four sites. The two sites in Massachusetts, USA: West Dennis (41.651918, -70.181307) and North Scituate (42.2375, - 70.780278), were populations also examined by Sanford et al. (2006). To examine how PLDs might vary over a larger spatial scale, we collected ovigerous females from two additional populations farther into the range core. In June 2022, we collected ovigerous females from Painter (37.519194, -75.7812) and Cushman’s Landing (37.17575, -75.9425), both located in Virginia, USA. These sites are located approximately 640 km south of the Massachusetts populations and are a similar straight-line distance from each other as the two Massachusetts populations. Because ovigerous females were not often seen on the surface of the marsh, we used trowels to excavate burrows and hand captured female crabs (Sanford et al. 2006). We only captured crabs with embryos that were gray in color, which is indicative of advanced stages of embryonic development (Phillips et al. 2010; Martins et al. 2020). We collected 30 crabs at each location and placed them in coolers for transport back to the Gloucester Marine Station (2019) and the University of Massachusetts Amherst (2022) for holding and experimentation. There were significant differences between carapace width between edge and core populations; however *M. pugnax* is known to follow Bergmann’s rule (Johnson et al. 2019), where populations in colder climates are larger than those in warmer climates. Carapace width was 13.2 ±0.357 mm (mean ± SEM) for Cushman’s Landing, VA, 14.0 ±1.58 mm for Painter, VA, 16.2 ±0.903 mm for West Dennis, MA, and 15.7 ±2.13 mm for Scituate, MA.

### Larval spawning and rearing

Once brought back to the lab, we placed ovigerous female crabs into 480 mL glass jars with 300 mL of seawater. We simulated tidal cycles by draining the jars of water for 10 hours during the day and refilling jars with seawater at night, which is often when *M. pugnax* release larvae in the wild (Morgan and Christy 1995). For the 2019 experiments we used 5 micron filtered natural seawater at the Gloucester Marine Station, diluted to a salinity of 27 ppt (Sanford et al. 2006) using reverse osmosis deionized (RODI) water (Milli Q Water Filtration system, Darmstadt, Germany). During the summer of 2022, we used artificial seawater mixed to a salinity of 27 ppt (Instant Ocean, Spectrum Brands, Blacksburg, VA) with RODI water at the UMass Amherst campus. Ovigerous females were held in individual culture jars in an 18□ water bath and maintained with daily 100% water changes until mothers released larvae.

To quantify the PLD, we reared larvae from mothers sourced from populations as described above. Several female crabs spawned during the first night of the spring tide, but the rest of the mothers exhibited variation in release dates, in contrast to Sanford et al. (2006). Therefore, we staggered the start of jars based on when the female crabs spawned (3 July 2019 - 19 July 2019 and 13 June 2022- 24 June 2022). Once larvae were released, we placed 40 larvae from each female crab into a new 480 mL glass jar with 300 mL of seawater with antibiotics (22.0 mg / L sodium penicillin G and 36.5 mg / L streptomycin; O’Connor and Judge 1997), which was the identical protocol conducted by Sanford et al. (2006). Jars were then placed into a temperature regulated water bath maintained at 18□ (±1□). We chose 18□ because this is the thermal threshold for larval development for *M. pugnax* and allows comparisons with historical measurements of PLD (Sanford et al. 2006). Crabs and larvae were exposed to an ambient and natural light photoperiod of approximately 15:9 hours (light:dark). During summer 2019, we cultured larvae from 6 mothers per population, while for 2022 we increased our sample size to 8 mothers per population. Larvae were fed live marine L-type rotifers, *Brachionus plicatilis* (Reed Mariculture, Campbell, California)*, ad libitum* with a target density of 40 rotifers per larva. Fiddler crabs undergo 5 stages of larval development (Sanford et al. 2006) and rotifers were replenished daily during zoeal stages 1-3. At zoeal stage 3, larvae were fed a mixture of marine rotifers and newly hatched *Artemia* sp., with the proportion of *Artemia* sp. increasing over larval development. During zoeal stages 4 and 5, larvae were fed mostly *Artemia* sp. with small amounts of marine rotifers to ensure that slower developing larvae were still able to access food. Consistent with Sanford et al. (2006), we conducted water changes every three days during summer 2019. However, because of a much lower survival rate (6%) in 2019 relative to Sanford et al. (2006) (survival >85%), we conducted daily water changes during summer 2022. Each jar was checked daily for the presence of metamorphosed larvae (i.e., megalopae), which were removed and either preserved in ethanol or used for the respirometry experiment (see below). We defined the pelagic larval duration (PLD) as the number of days from a larva’s spawning date to metamorphosis date. We ended the experiment when all the larvae metamorphosed or were no longer viable, defined here as when a larva could not swim off the bottom of the jar after 20 seconds. We used this approach because we generally observed healthy larvae to constantly swim and noted that larvae that are unable to move off the bottom of the jar were not likely to survive (Barley, *personal observation*).

### Pelagic larval duration data analysis

To determine if *M. pugnax* PLD varied spatially, we analyzed larval duration data collected in 2022 across the four edge and interior populations. Here, we used generalized linear mixed models (GLMMs) with a Poisson distribution to investigate the differences in PLD for this analysis and the temporal analysis described below. We used PLD as the response variable, population as a fixed effect, and mother as a random effect and we evaluated the fit of this model by comparing the idealized fit of a Poisson model to our model fit.

To determine whether PLDs of larvae from two edge populations had changed over time in association with warming ocean temperatures, we compared PLD data collected from Scituate and West Dennis, MA in 2019 and 2022 from this experiment with data collected in 2003 and 2004 by Sanford et al. (2006). We modeled Scituate and West Dennis separately and used PLD as the response variable, year as the fixed effect and mother as the random effect. We then completed a Tukey’s Honest Significant Differences test to compare the pairwise differences between the populations and we report the 95% confidence interval around each parameter estimate. Most of these models had singularity issues, likely because many larvae metamorphosed in the same jar on the same day, which decreased the variability in the data and causes the diagonals in the GLMM variance/covariance matrix to be zero or very close to zero. Therefore, in parallel to the GLMMs, we constructed a Cox survival regression model. We used metamorphosis (yes = 0, no = 1) as the binary response across time (days since spawning, PLD) and population as the predictor. We ran two models, one where the larvae were pooled within their population, and one with larvae pooled within the northern region (MA populations) and southern region (VA populations). We report the model with the highest log likelihood. For all models, we used ‘lme4’, ‘glmmTMB’, and ‘survival’ (Therneau and Grambsch 2000; Bates et al. 2015; Brooks et al. 2017; Therneau 2023) in R (version 4.3.0).

To determine if differences in PLD were associated with *in situ* sea-surface temperature (SST) data, we extracted environmental data from the NOAA Optimum Interpolation SST high-resolution data set which integrates satellite, ship, buoy, and float data at ¼° daily resolution (Huang et al. 2021). We downloaded raster data for each summer (May-September) for 2000-2022. Assuming that PLD is heritable, SST experienced previously by the parents during the summer of their larval development likely imposes selection on this trait. In addition, female *Minuca* spp. likely reproduce in their second or third year (Colby and Fonseca 1984). Therefore, we modeled the year the PLD data were collected with a lag of one, two, and three years in the SST data. For example, we used raster SST data from 2002, 2001, and 2000 as predictors for the PLD data collected in 2003 with a one, two, and three-year lag, respectively. Once we compiled the raster objects, we created a 100 km circular buffer, excluding terrestrial areas, around the Scituate and West Dennis, MA locations. We chose the buffer of 100 km because *M. pugnax* larvae export out of the estuary (Morgan 1990) into the coastal ocean and marine invertebrate larvae can often disperse this distance (Kinlan and Gaines 2003; Cowen and Sponaugle 2009). We then calculated SST summary statistics for each summer: average SST, average days above 18□, and average degree days. Average degree days were calculated using the temperature threshold for larval development (18□, Sanford et al. 2006) and subtracting this number from each mean daily temperature (Saco-Álvarez et al. 2010). We calculated a total of nine summary statistics for each year that PLD data were collected (2003, 2004, 2019, 2022): each of the above three summary statistics with a one-, two-, and three-year lag.

To model the effect of environmental conditions on PLD, we constructed generalized linear models with a Poisson error distribution, with PLD as the response variable, and year and summary statistic as the predictor variables. We also ran one model for each summary statistic for a total of eighteen models and compared all of these models using the corrected Akaike’s Information Criterion (AICc) for model selection. We considered the models with a delta AICc under 2 to be models that explained the variation in the PLD data well (Burnham and Anderson 2004). Model selection favored the inclusion of year in all models. In addition, we calculated the Variance Inflation Factor (VIF) for the models within two delta AICc units. The top model did not show evidence for collinearity, as the VIF values for SST statistic and year were both under 3 (Zuur et al. 2010). However, the other two models within two delta AICc units of the top model have VIF values of about 7, which indicates that SST and year were collinear. We did not further consider these collinear models.

### Larval respirometry

To examine whether variation in PLD across populations was correlated with metabolic rate, we measured oxygen consumption of a sub-sample of newly metamorphosed megalopae from the PLD experiments in 2022. Megalopae that metamorphosed within the previous 24-hours were placed into a separate container for holding with fresh seawater and fed newly hatched *Artemia*. When more than one larva metamorphosed on the same day from the same jar, megalopae were chosen at random for inclusion in the respirometry experiment. Megalopae not used in the respirometry experiment were preserved in 90% ethanol. We quantified metabolic rate using closed chamber static respirometry. Chambers were constructed with 7 mL glass scintillation vials and oxygen sensor spots (PreSens Precision Sensing GmbH, Model: SP-PSt3-NAU-D5-YOP-SA, Regensburg, Germany) affixed to the inside wall of the chambers. We then quantified oxygen content over time with a PreSens Fibox 4 Trace (PreSens Precision Sensing GmbH, Regensburg, Germany). Megalopae were placed into these vials without food and with fresh artificial seawater that was vigorously bubbled with air for one hour to ensure 100% oxygen saturation, making sure that there was no air left inside the vial. A control chamber blank with just seawater was run to quantify background respiration rates. Vials were then placed into a water bath kept at 18□ for the remainder of the experiment. We waited one hour to let the megalopae acclimate and then took one oxygen concentration reading per hour, for three hours the afternoon the experiment was started. The next morning, we took three more readings spaced one hour apart. After the experiment was completed, we preserved each megalopa in 90% ethanol. To calculate mass-specific metabolic rate, later we removed megalopae from ethanol, carefully dabbed them with a paper towel and allowed them to air dry for 15 minutes. We then weighed megalopae using a microbalance (model M2P, Sartorius, Göttingen, Germany). We calculated metabolism using the following equation, using the specific dynamic action term (Sp) of 0.014 kJ (Secor 2009):

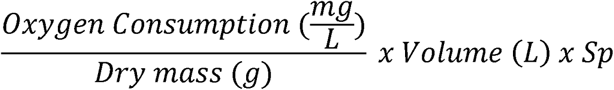

We determined oxygen consumption rates for the megalopae by applying linear regression models to the measured dissolved oxygen in each chamber for the duration of the experiment. We recorded the slope of the best-fit model as the oxygen consumption for each individual megalopa and background oxygen consumption rates. We then subtracted background respiration for a given trial from the observed metabolic rate of all megalopae from the same trial. To compile and compute respiration rates from the oxygen concentration data, we used the respR package in R (Harianto et al. 2019). To analyze the differences in metabolic rate among the *M. pugnax* populations, we used generalized linear mixed models (GLMMs) with a Gaussian error distribution. We used metabolic rate as the dependent variable, population as a fixed effect, and day of the experiment was included as a random effect.

## Results

In 2019, 16/240 larvae survived to metamorphosis from Scituate, MA and 31/240 from West Dennis, MA, and there was no significant difference in PLD between these populations (p = 0.88, z = -0.148, df = 45). In 2022, 443 larvae (out of 1,280) survived to metamorphose into megalopae. Pooled across all sites, the average PLD at 18□ was 35.3 days, and ranged from 20 to 54 days. The northern, edge populations (Scituate and West Dennis, MA) had significantly faster larval development times than the southern, core populations (Cushman’s Landing and Painter, VA, Table 1, Table 2, Fig. 2). Scituate and West Dennis had PLDs that were 3.2 (±1.1) and 2.4 (±1.3) days faster than Cushman’s Landing, VA, respectively (Scituate: p<0.0001; West Dennis: p<0.0001). Development rates for Scituate and West Dennis were 2.4 (±1.4) and 1.5 (±1.4) days faster than Painter, VA, respectively (Scituate: p<0.0001; West Dennis: p=0.02). Development rates did not differ between the two sites in Massachusetts (p=0.39) or between the two sites in Virginia (p=0.19). The Cox survival regression reflects the same pattern, where Scituate and West Dennis larvae developed significantly faster than Cushman’s Landing larvae (p>0.000 and p=0.0002, respectively; Table S1, Figure S1). The regression that had larvae pooled into the four populations had a higher log likelihood and is therefore the model we are reporting here (LL: -2237.9 versus -2240.9). We ran the GLMM to account for the effect of mother on PLD, however this effect was not significant and the parameters from the GLMM and the GLM without the mother effect were identical. We did not find any significant differences across populations in the metabolic rates of *M. pugnax* megalopae (Fig. 5; Table S2).

**Figure 2.**
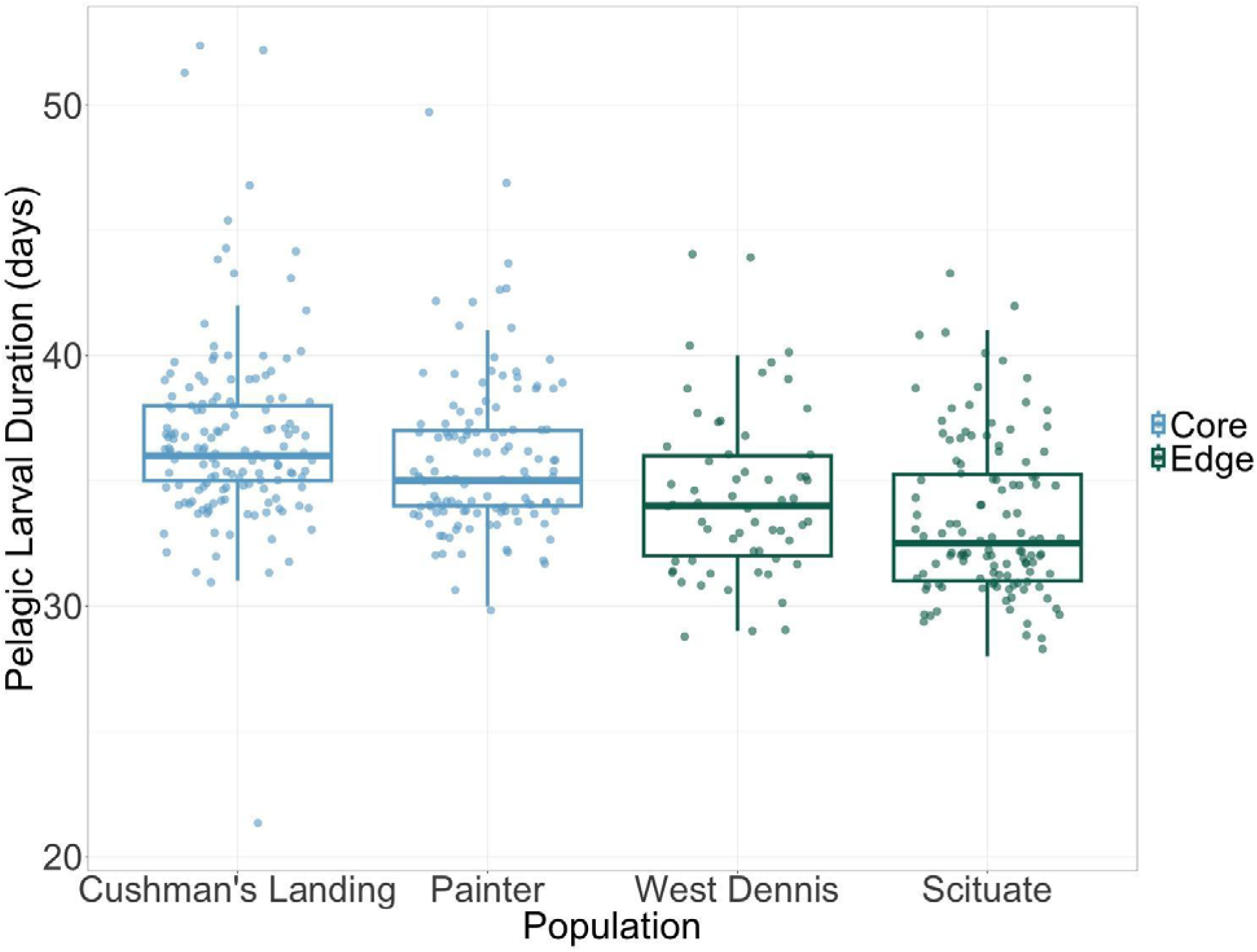
Box plots and overlaid points of pelagic larval duration (PLD) reared at 18□ by each of the *Minuca pugnax* populations in the 2022 larval development experiment. Sites are ordered left to right (core to edge). Each data point represents the PLD for an individual larva, pooled across all jars/mothers, and jittered to reduce the overlap of data points. Boxplots overlaid on top of the data points represent quartiles for each population.

**Table 1.** GLMM parameter estimates for the mud fiddler crab 2022 experiment. The model was run with pelagic larval duration (PLD) as the response, population as the predictor, and jar/mother as the random effect. Asterisk (*) denotes intercept.

| Term | Estimate | SE | Z value | P value |
| --- | --- | --- | --- | --- |
| Core 1<br>(Cushman's, VA)* | 3.603 | 0.0136 | 263.8 | <0.001 |
| Core 2<br>(Painter, VA) | -0.0228 | 0.0204 | -1.115 | 0.2647 |
| Edge 1<br>(West Dennis, MA) | -0.0667 | 0.0259 | -2.573 | 0.0101 |
| Edge 2<br>(Scituate, MA) | -0.0915 | 0.0211 | -4.342 | <0.001 |

**Table 2.** Post-hoc comparisons of pelagic larval durations (PLD) between populations of mud fiddler crabs for the model in Table 1. CI = confidence interval.

| Comparison | Difference | Lower CI | Upper CI | P value |
| --- | --- | --- | --- | --- |
| Edge 1 – Core 1 | -2.369 | -3.704 | -1.034 | <0.001 |
| Edge 2 – Core 1 | -3.211 | -4.293 | -2.128 | <0.001 |
| Edge 1 – Core 2 | 1.542 | 0.168 | 2.917 | <0.001 |
| Edge 2 – Core 2 | -2.384 | -3.515 | -1.253 | <0.001 |
| Edge 1 – Edge 2 | -0.841 | -2.226 | 0.543 | 0.398 |
| Core 1 – Core 2 | -0.827 | -1.897 | 0.244 | 0.193 |

We modeled the PLD data from 2003, 2004, 2019, and 2022 for each population separately and found significant decreases in PLD across time for both Scituate and West Dennis (Table 3, Fig. 3). Larvae from Scituate, MA developed 2.3 (±2.0) days faster in 2022 than in 2003 (Table 3) at 18□, and larvae from West Dennis developed 5.3 (±2.0) days faster in 2022 than in 2003 (Table 3). For the remote-sensed SST analysis, the additive model that included average days above 18□ with a two-year lag was the best supported model (Table 4), which indicated that with an increasing average number of days above 18□, PLD at 18□ significantly decreased over time from 2003 to 2022 by 4.94 days and 2.34 days in West Dennis and Scituate, respectively (Table 5, Fig. 4). PLDs in 2004 were significantly faster than those in 2003 (p=0.004) and PLDs in 2019 did not differ significantly from those in 2003 (p=0.074).

**Figure 3.**
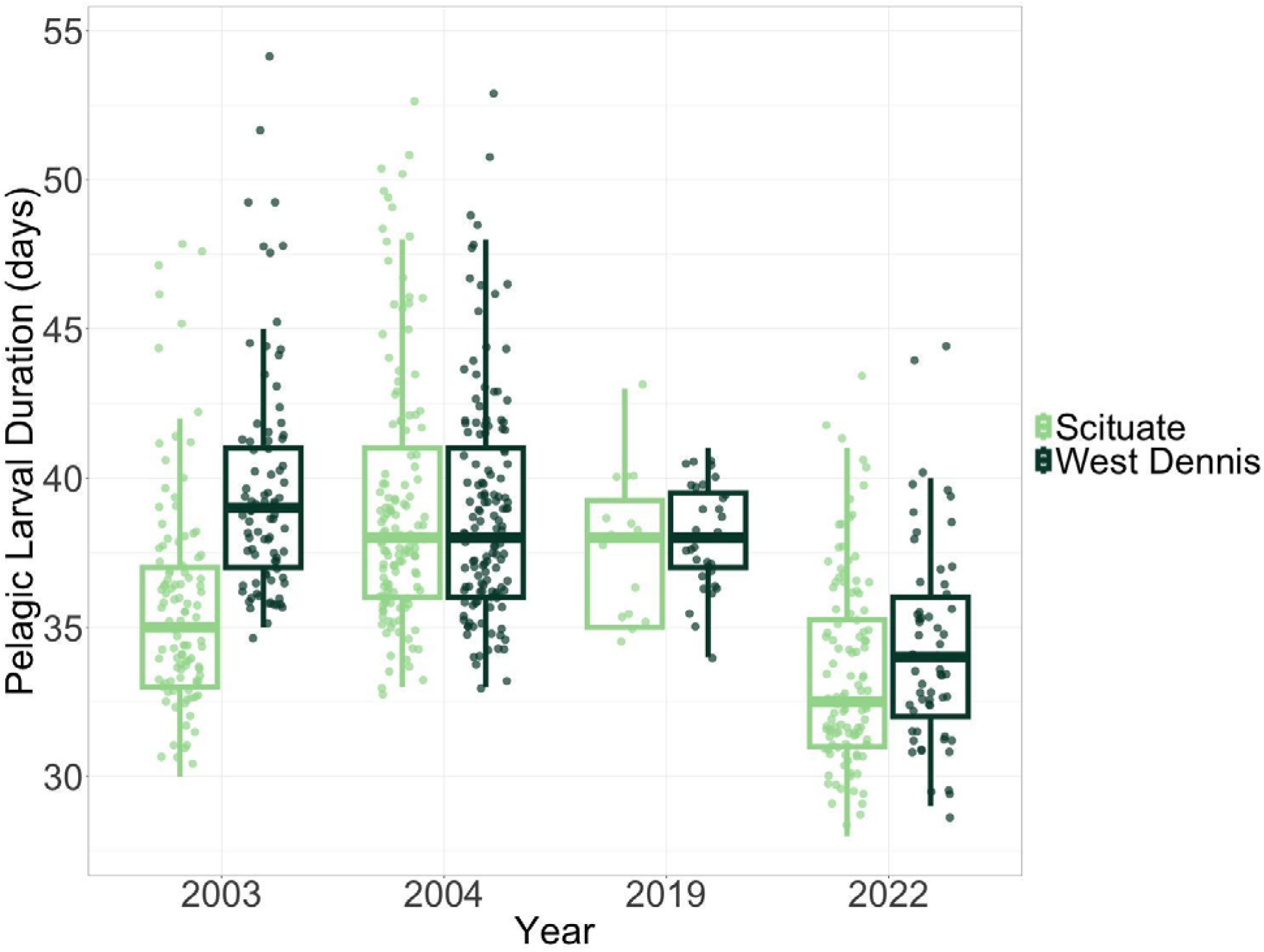
Box plots and overlaid points showing the larval duration of *Minuca pugnax* reared at 18°C, across years and populations. Both Scituate and West Dennis are edge populations. The 2003 and 2004 data were collected by Sanford et al. (2006). Each data point represents the PLD for an individual larva, pooled across all mothers, and jittered to reduce the overlap of data points.

**Figure 4.**
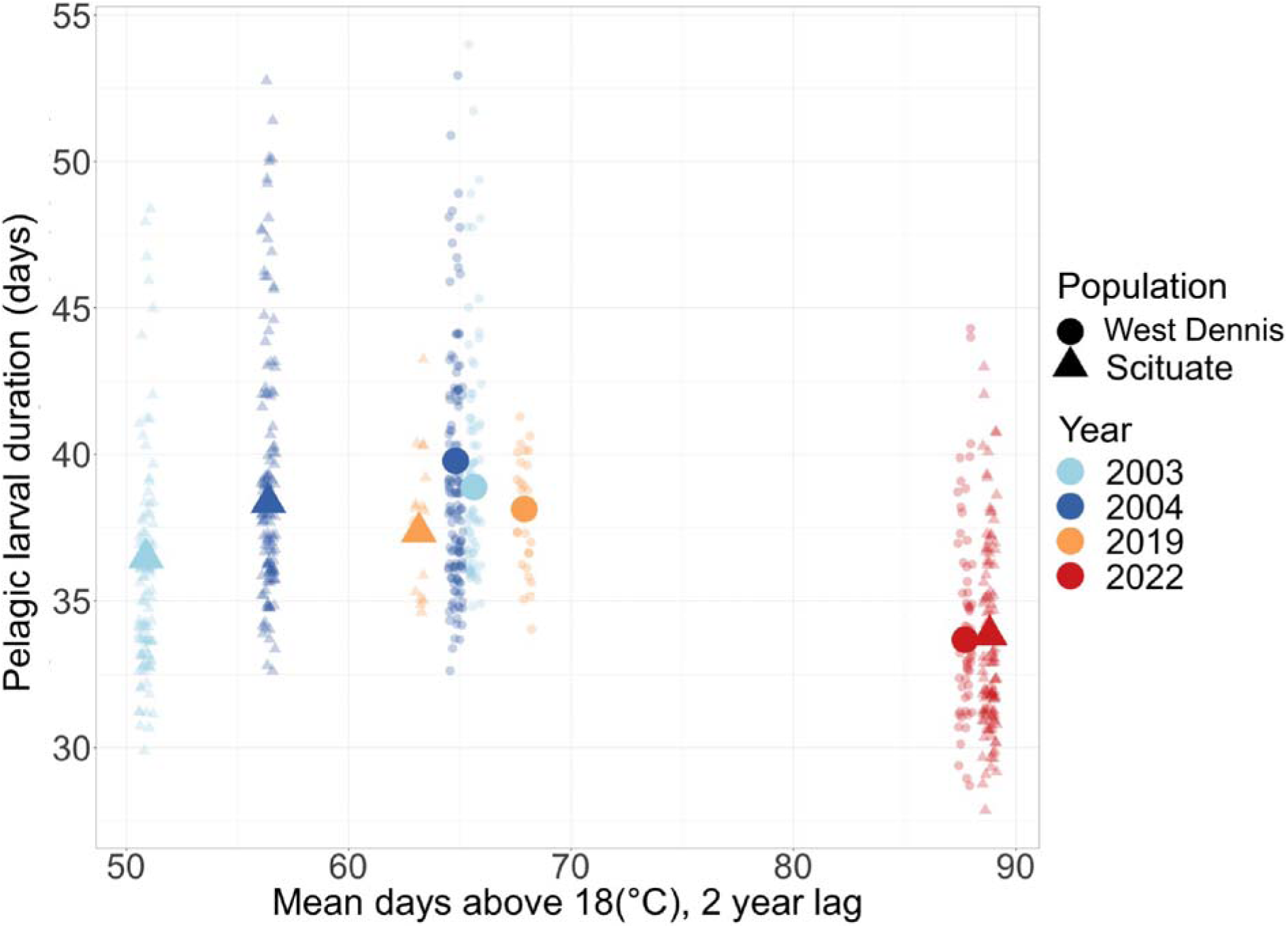
Pelagic larval duration data for the edge populations as a function of the average days above 18□ with a two-year lag. Bolder points are model predictions for each population and year. Average days above 18□ for each year and population were calculated using raster OISST data from NOAA (Huang et al. 2021).

**Figure 5.**
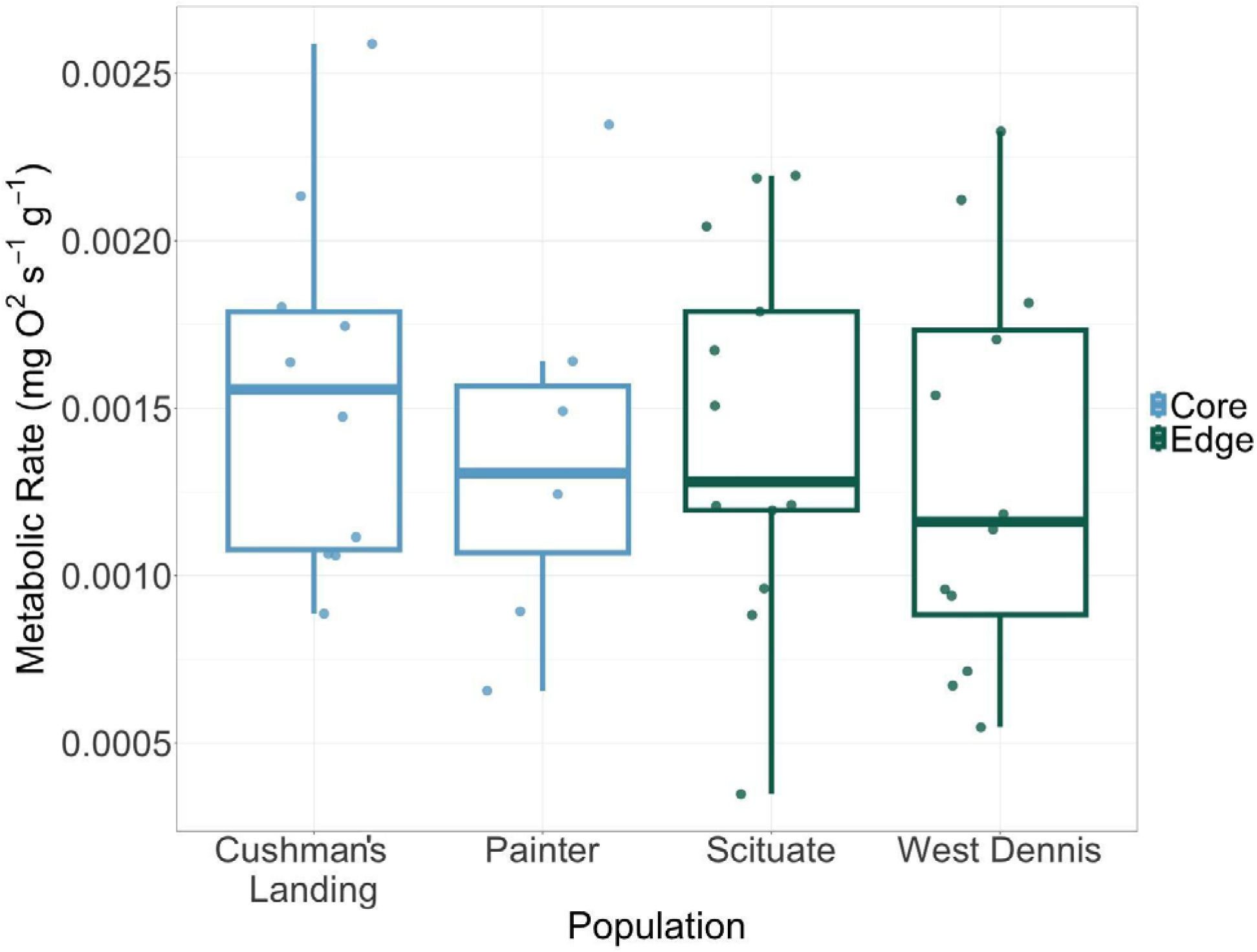
Mass-specific metabolic rate of *Minuca pugnax* megalopae for range core and edge populations in the 2022 experiment. Each data point represents the mass-specific rate at which each megalopa consumed oxygen in a closed chamber at 18□.

**Table 3.**
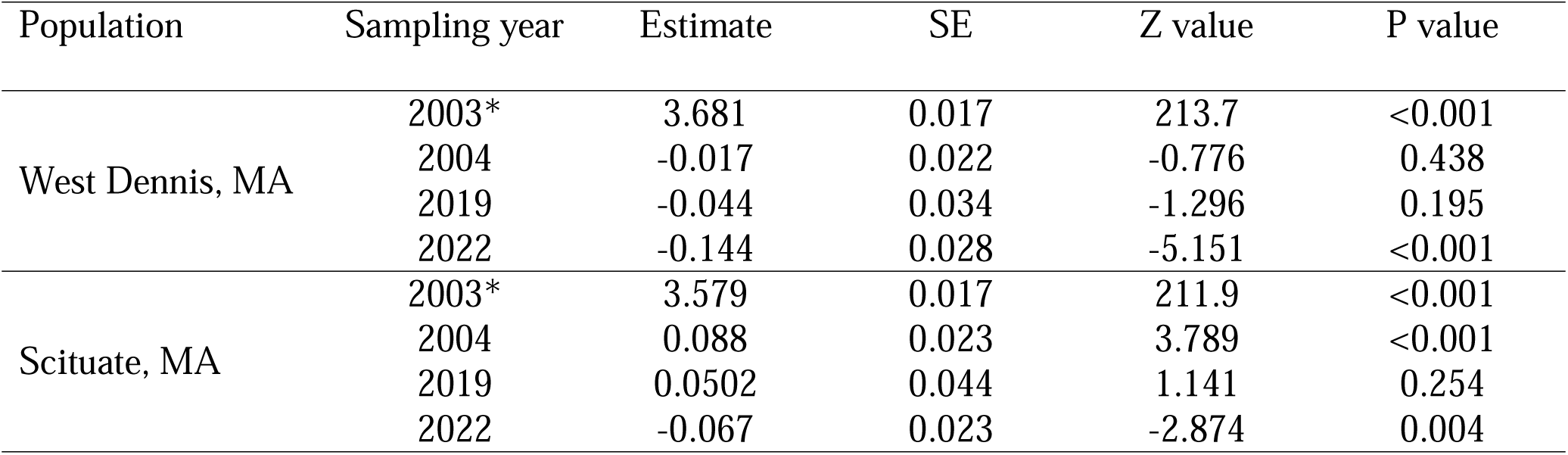
GLMM parameter estimates for Scituate and West Dennis, MA for quantifying PLD across time. The model was run with pelagic larval duration (PLD) as the response, sampling year as the predictor, and jar/mother as the random effect. Asterisk (*) denotes intercept.

**Table 4.** Model selection table for models for the sea surface temperature (SST) analysis with Variance Inflation Factor (VIF) values lower than three. Additive models with the year parameter are marked. LL= Log Likelihood, Wt= Weight, AICc= Corrected Akaike Information Criterion.

| Model name | K | AICc | Delta AICc | AICc Wt | Cum Wt | LL |
| --- | --- | --- | --- | --- | --- | --- |
| Average days above 18□, 2 yr. lag (additive, year) | 5 | 3999.3 | 0.00 | 0.97 | 0.97 | -1994.6 |
| Average days above 18□, 3 yr. lag (additive, year) | 5 | 4006.0 | 6.68 | 0.03 | 1.00 | -1998.0 |
| Average summer SST, 2 yr. lag | 5 | 4020.8 | 21.52 | 0.00 | 1.00 | -2008.4 |
| Average degree days, 2 yr. lag | 5 | 4020.8 | 21.52 | 0.00 | 1.00 | -2008.4 |
| Average summer SST, 1 yr. lag | 5 | 4025.9 | 26.58 | 0.00 | 1.00 | -2010.9 |
| Average degree days, 1 yr. lag | 5 | 4025.9 | 26.58 | 0.00 | 1.00 | -2010.9 |
| Average days above 18□, 1 yr. lag | 5 | 4030.7 | 31.39 | 0.00 | 1.00 | -2013.3 |
| Average days above 18□, 2 yr. lag | 5 | 4043.4 | 44.15 | 0.00 | 1.00 | -2019.7 |
| Average degree days, 3 yr. lag | 5 | 4072.3 | 73.06 | 0.00 | 1.00 | -2034.2 |
| Average summer SST, 3 yr. lag | 5 | 4072.3 | 73.06 | 0.00 | 1.00 | -2034.2 |
| Average days above 18□, 3 yr. lag | 5 | 4072.8 | 73.47 | 0.00 | 1.00 | -2034.4 |

**Table 5.** Model summary table for the top SST model; this is an additive model with year and average days above 18□ with a two-year lag.

| Term | Estimate | SE | Z value | P value |
| --- | --- | --- | --- | --- |
| Year, 2003 | 3.376 | 0.076 | 44.24 | <0.001 |
| Average days above 18□,<br>2 yr. lag | 0.004 | 0.001 | 3.332 | <0.001 |
| Year, 2004 | 0.026 | 0.016 | 1.671 | 0.095 |
| Year, 2019 | 0.026 | 0.015 | -1.020 | 0.308 |
| Year, 2022 | -0.240 | 0.044 | -5.471 | <0.001 |

## Discussion

The evolutionary mechanisms underlying range shifts are poorly understood, particularly in marine systems that exhibit rates of spread that can be up to an order of magnitude faster than in terrestrial ecosystems (Parmesan and Yohe 2003; Sorte et al. 2010; Chen et al. 2011; Poloczanska et al. 2013). In systems with prevailing equatorward flows, theory suggests that poleward range shifts are favored in species with rapid larval development that allows propagules to disperse poleward during episodic and short-lived current reversals (Byers and Pringle 2006). Strikingly, our common garden experiments support this hypothesis; larvae from edge populations had significantly shorter larval durations (1.5-3.2 days shorter on average) as compared to larvae from core populations, despite being reared at the same temperature in the laboratory. In addition, our lab trials across time revealed evidence for accelerated larval development in 2022 (but not 2019), highlighting the potential for change in PLD over time.

### Spatial sorting

Our results contribute to a growing body of research on the role of spatial sorting in promoting range expansions in terrestrial systems. For example, spatial sorting influences a variety of dispersive traits including boldness behavior in spiders (Chuang and Riechert 2022), morphology in an invasive bird (Berthouly-Salazar et al. 2012), seed weight in plants (Latron et al. 2022), and higher frequency of long-winged morphs of bush crickets (Simmons and Thomas 2004). These studies found evidence for spatial sorting on scales between 100 and 700 km. However, marine organisms have greater dispersal on average when compared to terrestrial species (Kinlan and Gaines 2003; Sorte et al. 2010) and therefore the scale of spatial sorting is likely larger than in terrestrial systems. Sanford et al. (2006) found variation consistent with spatial sorting in *M. pugnax* populations sampled on the scale of 100 km. In the present study, we provide support for the spatial sorting hypothesis on the scale of 715 km.

Under spatial sorting, populations near the range edge should accumulate high dispersers; however, as a range expansion occurs and previous edge populations become part of the range core, the frequency of higher dispersing individuals should decrease (Simmons and Thomas 2004). In our system, if former edge populations have become part of the range core, we would expect average PLDs to increase, as high dispersing individuals with shorter PLDs no longer collect at high frequencies in these populations. Instead, our results show that there was a dramatic decrease in PLD over time in Scituate and West Dennis. We note however, that the PLD in 2019 was not different from prior years, which may have been driven by a lower sample size, possibly combined with strong interannual variation, which was also observed in 2003 and 2004. In addition, while we might theorize that under spatial sorting PLDs should increase over time, this is not the only process that is likely affecting PLDs in this system. The Gulf of Maine, and the larger oceans, are warming rapidly, possibly generating a selective environment that favors faster development and subsequently shorter PLDs. The mechanisms producing this pattern remain unclear but likely involve tradeoffs whereby faster development comes at a cost (Arietta and Skelly 2021).

Given that *M. pugnax* has been observed poleward of our edge sites as far north as central Maine, USA (Martínez-Soto and Johnson 2020), it is likely that spatial sorting in mud crabs operates across a larger spatial scale, where the “edge” encompasses our two edge sites as well as the realized range edge in Maine. This is supported by the large dispersal potential of mud fiddler crabs (30+ day PLD) and coastal oceanography in the Gulf of Maine which can transport propagules long distances and promote population connectivity at large spatial scales (Xue et al. 2008; Li et al. 2014). It is possible that the Scituate and West Dennis populations are still a part of the range edge and are acting as source populations to the farthest poleward population. In this case, larvae that were spawned south of Boston, MA in more established populations may disperse poleward to sink populations that are not yet able to sustain larval self-recruitment. For species like *M. pugnax* that disperse in currents to sustain populations upstream, individuals that settle have a high likelihood of having shortened PLDs because this increases the chance that they would settle in the most poleward populations (Sanford et al. 2006; Allgayer et al. 2021). While it is well understood that ocean currents generally play a large role in marine metapopulation dynamics (Watson et al. 2012; Allgayer et al. 2021), it is not well understood how metapopulation dynamics and currents affect spatial sorting in the leading edge of a range expanding species.

### Countergradient variation

Countergradient variation arises from selection imposed by environmental conditions that result in traits that are in opposition to environmental influences (Conover and Present 1990; Conover and Schultz 1995; Arendt and Wilson 1999). In our system, spatial sorting and countergradient variation would present similarly, with shorter PLDs in the colder, edge populations. To discriminate between these hypotheses, we evaluated the potential for CnGV to operate over time with *in situ* SST data. The GOM warmed from 2000 to 2022 where in Boston Harbor, the average days above 18□, a critical threshold for larval development, nearly doubled to about 80 days (Fig. 1). The countergradient variation hypothesis predicts an increase in PLD due to the release from the selective pressure of cold, short, growing seasons because theory suggests that such selection can occur over both space and time (Conover et al. 2009). In contrast, we found that PLD decreased from 2003 to 2022, suggesting that CnGV is possibly not operating in this system. However, it should be noted that *M. pugnax* can produce more than one brood per spawning season; larvae that are spawned later in the season in higher latitudes are more time constrained than those spawned earlier in the season (i.e., intra-annual variation in growth rates; Dmitriew 2011). Additional tests examining intra-annual variation in PLD (i.e., early versus late season spawning crabs) could further resolve this possibility. In addition, shorter PLDs might be selected for because extended larval development in cooler waters leaves the individual susceptible to predation; therefore, spending less time as planktonic larvae would be advantageous (Thorson 1950; Chambers and Leggett 1987; Takasuka et al. 2007; Robert et al. 2023). However, while this might explain the variation in PLD across space, it cannot explain the decrease in PLD that we observed in the edge populations across time. While these results cannot completely disentangle the spatial sorting and the countergradient variation hypotheses, taken together we interpret our data to be most consistent with spatial sorting.

### Metabolic rate

As a potential mechanism underlying accelerated larval development in edge populations, we examined larval metabolic rate across populations from our common garden experiments. We expected that faster development rates in colder, edge populations would be accompanied by higher metabolic rates (metabolic compensation hypothesis, Conover et al. 2009). Although larvae from colder, edge populations had faster larval development rates, we did not observe differences in standard metabolic rate in megalopae. This is consistent with earlier reports of no differentiation in metabolic rate of *M. pugnax* megalopae sourced from New York (higher latitude) as compared to those from North Carolina (lower latitude) when reared at lower temperatures (Vernberg and Gostlow 1966). Since we examined respirometry of megalopae, it is possible that metabolic rates of earlier larval stages are greater for edge as compared to core populations. However, prior work has also shown that there is not population differentiation for first-stage zoea (Vernberg and Gostlow 1966). Instead, differences in larval duration might be associated with differences in energy allocation. Marine larvae may allocate energy to different biological processes such as growth, protein synthesis, or lipid production based on endogenous growth rate or environmental stress (Pace et al. 2006; Frieder et al. 2018; DellaTorre and Manahan 2023). One possibility is that *M. pugnax* larvae have similar overall metabolic rates across populations, but individuals from edge populations allocate more energy to growth than energy reserves. In addition, maternal energy investment could vary across populations, which could affect growth (Marshall and Keough 2007) and larval development rates. For example, mothers from edge populations might invest in fewer, larger larvae than those from core populations that are able to develop faster.

## Conclusions

Because most marine taxa have a planktonic larval stage, our results have implications for understanding dispersal and range shifts more broadly. Globally, many important and highly productive marine systems are characterized by equatorward flows. Thus, dispersing poleward against an equatorward prevailing current is likely common, which could explain among and within taxa variation in direction and rates of spread (Sunday et al. 2012; Sanford et al. 2019). Our work highlights how ocean warming can facilitate spread by creating a more favorable environment for the completion of larval development or by shortening the time in the plankton, allowing propagules to take advantage of variable flows. In this case, spatial sorting may be a more general response for benthic marine species but needs further investigation. For example, many species with pelagic larval stages exhibit further complexities, such as swimming behaviors that can alter their dispersal potential. Interestingly, global analyses have revealed that the mismatch between current flow and direction of climate warming is more common in species with passive dispersal compared to those with more active dispersal mechanisms (García Molinos et al. 2017). However, in species that are expanding their range in alignment with prevailing ocean currents, warming may actually slow range expansion (O’Connor et al. 2007; Shanks 2009) and decrease population connectivity (Marshall and Alvarez-Noriega 2020). Our study adds to a growing body of research investigating the effects of microevolutionary processes, such as local adaptation and spatial sorting, that affect range limits and expansions. Mechanistic studies that reveal how ecological and evolutionary processes drive range shifts can help clarify our understanding of climate impacts on species redistribution globally.

