## Supplemental Materials for "Spatial sorting and accelerated larval development in a range expanding fiddler crab"

**Supplemental Table 1.** Summary table for the Cox survival regression constructed from the 2022 larval Pelagic Larval Duration (PLD) data, where larvae are pooled across our four populations. The intercept for this model is the Cushman’s landing, VA population.

| Term | Coefficient | Exp(Coefficient) | Standard Error | Z value | P value |
| --- | --- | --- | --- | --- | --- |
| Population: Dennis | 0.5649 | 1.7592 | 0.1545 | 3.657 | 0.000255 |
| Population: Painter | 0.2248 | 1.2520 | 0.1238 | 1.816 | 0.069412 |
| Population: Scituate | 0.8265 | 2.2852 | 0.1266 | 6.530 | <0.0001 |

**Supplemental Table 2.** Summary table for Tukey’s Honest Significant Different post-hoc comparison of the GLMM model comparing the metabolic rates of megalopae across four populations (Scituate, MA, Dennis, MA, Painter, VA, and Cushman’s Landing, VA). CI= Confidence Interval

| Population Comparison | Difference | Lower CI | Upper CI | P value |
| --- | --- | --- | --- | --- |
| Dennis-Cushman | 2.456069e-04 | -0.0002678140 | 0.0007590279 | 0.5731606 |
| Painter-Cushman | 1.824061e-04 | -0.0004085134 | 0.0007733257 | 0.8374519 |
| Scituate-Cushman | 1.292685e-04 | -0.0003750967 | 0.0006336338 | 0.8989270 |
| Painter-Dennis | -6.320082e-05 | -0.0006334833 | 0.0005070817 | 0.9904617 |
| Scituate-Dennis | -1.163384e-04 | -0.0005963598 | 0.0003636830 | 0.9128330 |
| Scituate-Painter | -5.313758e-05 | -0.0006152811 | 0.0005090059 | 0.9940221 |

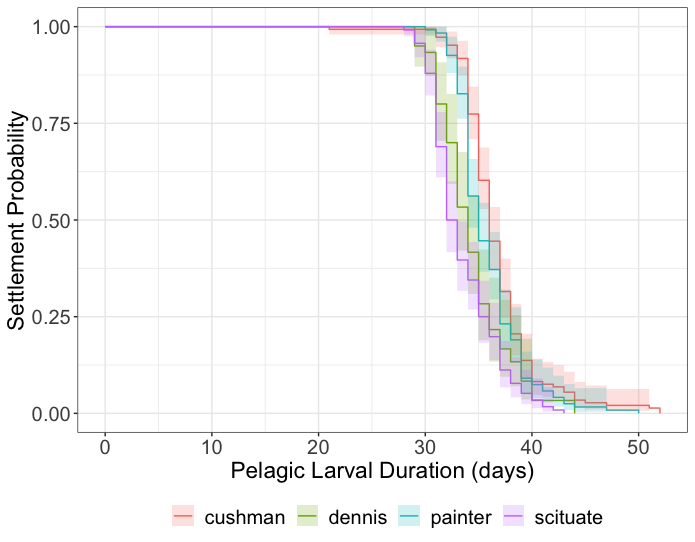

**Supplemental Figure 1.** Logistic curves depicting the Cox survival regression of the 2022 larval Pelagic Larval Duration (PLD) data. Settlement probability denotes the probability on any given day that a larva would metamorphose into a megalopae (or, commonly known as ‘settlement’). Color denotes the population that the larva came from; Cushman and Painter, VA are our southern (equatorward) populations, and Dennis and Scituate and our northern (poleward) populations.
